# Engaging working memory following skill reactivation has implications for interlimb skill generalization

**DOI:** 10.64898/2026.05.11.724282

**Authors:** Rahul Pal, Goldy Yadav, Neeraj Kumar

**Author notes:** **Data Availability:** The datasets used and/or analysed during the current study are available on request from the authors. Equally contributed to the work (Rahul Pal, Goldy Yadav).

## Abstract

Interlimb skill generalization, defined as the transfer of a newly learned skill from the trained to the untrained limb, represents a fundamental aspect of human motor behavior with significant implications for rehabilitation and athletic training. Skill generalization is influenced by processes that drive learning and interact with the newly acquired memory. For instance, in our recent work, we reported that performing a secondary, cognitively demanding task immediately after a short skill-training session impaired skill generalization when the untrained arm was tested 24-hour later. This suggests that working memory (WM) interacts with the early stage of skill memory consolidation processes and thereby impacts skill generalization. Motivated by this finding, in the current study, we investigate how WM interacts with reactivated skill memory and its subsequent impact on skill generalization, tested 24 or 48-hour post skill training. We recruited right-handed young participants (n=95) who performed a fast, accurate reaching task with their dominant right arm during a short training session (50 trials) on Day-1. After 24-hour on Day-2, depending on the group type, participants had a brief skill reactivation session (10 trials or no reactivation) and then performed the WM task (or a control task) with their right arm. Interlimb generalization to the untrained left arm was assessed either immediately after the WM/control task on Day-2 or after a 24-hour gap on Day-3. We found that, engaging in the WM task (compared to the control task) after skill reactivation on Day-2 enhanced immediate generalization. Conversely, when generalization was tested 24-hour later on Day-3, the same WM engagement impaired skill generalization. These findings demonstrate that WM engagement during the post-reactivation phase has a time-dependent influence on interlimb generalization. WM can facilitate immediate generalization, possibly by sustaining neural processes that promote skill memory generalization across effectors. However, when a 24-hour’s time gap is introduced, generalization is disrupted following WM engagement, possibly because of interference between underlying neural processes involved in WM and reactivation-induced (re)consolidation of the skill memory. This study highlights the delicate interplay among WM, motor memory reactivation dynamics, and skill generalization and suggests a time-dependent interplay of neural processes critical for optimizing outcomes in motor learning and clinical rehabilitation protocols.

**NEW & NOTEWORTHY:** This study reveals that working memory engagement following skill memory reactivation exerts a time-dependent dual effect on interlimb generalization. When tested immediately, WM engagement enhances generalization by prolonging the labile state of reactivated memory, which is needed to transfer abstract/effector-independent features to the untrained arm. When tested 24-hour after a delay, the same intervention impairs generalization by possibly engaging reconsolidation processes that interfere with effector-independent features. These findings establish a unified framework in which the temporal dynamics of the reconsolidation window determine whether a cognitive intervention strengthens or weakens a motor memory, with implications for understanding memory plasticity and for designing effective rehabilitation protocols.

## INTRODUCTION

Interlimb generalization (also known as intermanual transfer or cross education) is considered a key aspect of human skill behavior and motor rehabilitation in which learning with one limb incurs performance improvements to the untrained limb. Interlimb generalization is typically measured (Yadav & Mutha, 2020) by comparing the naïve (initial) performances of the trained and the untrained arms on a given task either performed on the same day or on a different day. Our past work on interlimb generalization shows that newly learned skilled movements can transfer to the untrained arm immediately following training on the same day (Day-1)-from the dominant right arm to the non-dominant left arm and vice versa (Yadav & Mutha, 2020). In our recent studies (Yadav, Pal, et al., 2025; Yadav, Vassiliadis, et al., 2025) we noted that such newly learned skills can generalize to the untrained non-dominant arm even when tested 24-hour following training (Day-2) - a time period which normally allows freshly acquired skill memory to stabilize/consolidate. Interestingly, such interlimb generalization at 24-hour was found to be robust in the case of a long skill training session and could not be interfered by engaging in secondary tasks post training (Yadav, Pal, et al., 2025). However, performing a secondary task that is cognitively demanding (such as a working memory (WM) task) after a short session of skill training led to an impairment in interlimb generalization when the untrained arm was tested after 24-hour. These findings imply that WM task can interfere with interlimb generalization on Day-2 of a newly formed motor skill memory resulting from a brief skill training session on Day-1.

Additionally, it is known that newly formed motor memories are initially fragile and highly susceptible to interference or modifications from a secondary task thus impacting the follow-up performance (Brashers-Krug et al., 1996; Krakauer et al., 2005; Sing & Smith, 2010). For instance, a short/early phase of motor skill training and WM processing may compete for shared neural resources (such as the DLPFC-see Curtis & D’Esposito, 2003; D’Esposito & Postle, 2015; Miller & Cohen, 2001). Evidence from interference studies supports this view-introducing a cognitively demanding WM task immediately after skill training can disrupt initial skill consolidation, particularly when training is brief (Brown & Robertson, 2007; Keisler & Shadmehr, 2010; Seidler et al., 2012). While the effect of WM task on initial skill memory consolidation following Day-1 training is reported, the impact of WM engagement following a brief session of skill training on Day-2 is unknown. Such brief session of skill training on Day-2 following a time gap of several hours after initial training can be considered as a case of “reactivation” (Amar-Halpert et al., 2017; Herszage et al., 2023; Herszage & Censor, 2017). Research on memory reactivation suggests that subsequent brief exposure to the same task after training can reactivate the initial memory such that the stabilized/consolidated memory becomes unstable/fragile and can either strengthen or weaken depending on what happens in the post reactivation time window (Alberini & LeDoux, 2013; Birbaumer, 2010; Censor et al., 2010; Nader et al., 2000; Nader & Hardt, 2009; De Beukelaar et al., 2016; Herszage et al. 2021).

Thus, we wondered whether WM engagement after a brief session of skill reactivation would (case-A) destabilize the skill memory representations through resource competition between the skill memory and WM processing, or (case-B) strengthen the reactivated skill memory through overlapping neural processing. In the present study, we used this conceptual framework to investigate how WM engagement on Day-2 after skill reactivation will influence the follow up interlimb skill generalization to the untrained arm (tested on Day-2 or Day-3). Building on our prior findings (Yadav, Pal, et al., 2025), in which WM engagement on Day-1 disrupted skill generalization to the opposite limb, we hypothesized that WM engagement after a brief episode of skill reactivation on day-2 would interfere with further generalization to the opposite limb tested on Day-2 or Day-3 as outlined in Case-A. To test this hypothesis, we designed a series of three experiments involving right-handed young individuals (n=95) who received a short skill training on Day-1 (similar to that Yadav, Pal, et al., 2025; Yadav, Vassiliadis, et al., 2025). On Day-2, there was a session of brief (or no) memory reactivation, followed by WM or a control memory task. All participants were finally tested for interlimb generalization either on Day-2 or Day-3. We found perplexing results-WM engagement after reactivation led to enhanced interlimb generalization (contrary to our hypothesis, thus favoring case-B) on Day-2, whereas impaired generalization (in line with our hypothesis favoring case-A) on Day-3. These findings suggest that the interfering effect of WM on skill generalization (as observed in our previous study of (Yadav, Pal, et al., 2025) and in the current study when tested on Day-3) can become surprisingly facilitatory when there is no time gap between reactivation, WM task and the interlimb generalization session.

## METHODS

### Participants

Ninety-six young healthy adults completed all sessions of this study that spanned over 2-3 days (see Study Design, Figure 1) and 95 (Age range: 18-38 years old; Mean age: 22.30 ± 3.07; 34 Females) were included in the final data analyses (see Data Analyses section later). To estimate the required sample size, an a priori power analysis was conducted using G*Power 3.1 (Faul et al., 2009) for a repeated-measures design with a within–between interaction. Based on prior work (Yadav, Pal, et al., 2025), a moderate effect size (f = 0.25), significance level (α = 0.05), and statistical power (1 − β = 0.80) were used as input parameters. The analysis indicated that a minimum total sample size of 60 participants would be required to detect the expected effects. All participants were right-handed, as determined by the Edinburgh Handedness Inventory (Oldfield, 1971) and were naive to the experimental procedures. Written informed consent was obtained from all participants prior to the study, which the Institute Ethics Committee approved. None of the participants reported any history of neurological, medical, or psychiatric disorders and had normal or corrected-to-normal vision. All participants received monetary compensation for their participation.

**Figure 1:**
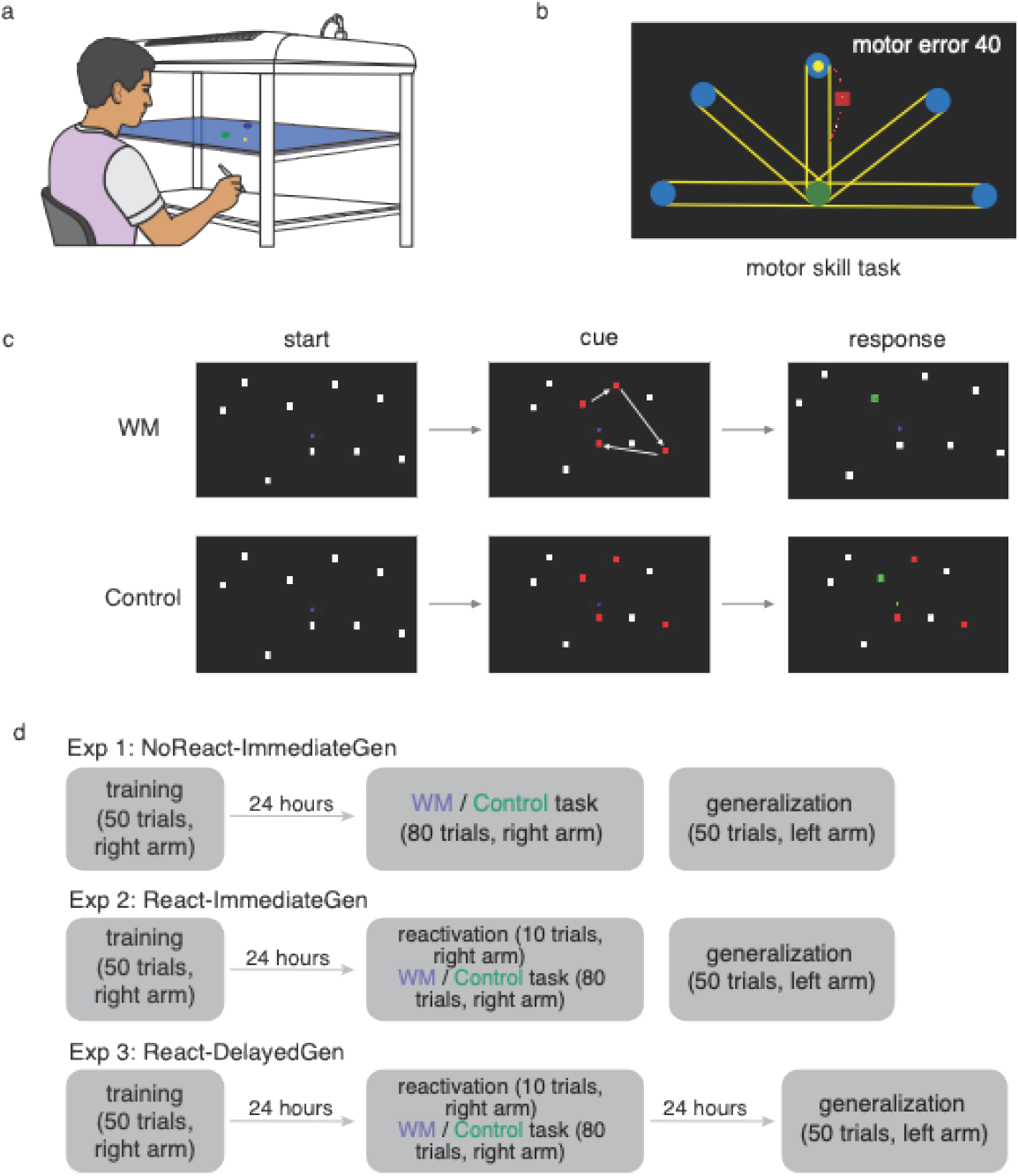
(a) Experimental setup with computer screen (top), reflective one-way mirror (middle), digitizing tablet (bottom) on which participants performed all the tasks. (b) Motor skill task with a representative movement from start circle (green color) to one of the five target circles (blue color) at a distance of 150 mm, a parallel yellow line connecting the start circle with the target circle. Note that only one target appears on each trial. Distance between the red square (indicating hand position at 650 ms of the executed movement) and target circle is considered temporal error. Red line indicating hand movement outside the parallel yellow path is considered spatial error. A composite sum of these two errors is indicated as Motor Error. Participants are instructed to reduce this Motor Error during training. (**c**) WM task (modified version of corsi-block task) indicating start (presentation of nine white squares with blue start circle and yellow dot indicating hand position), cue (in this representation four out of nine squares changed from white to red in a sequence). Note that the color change was sequential i.e., white squares flashed in red for 750 ms which was to be recalled later testing the working memory. However, for the control task the red squares were present on the screen for the entire trial duration, and the response screen (during which participants moved to touch the squares that turned red). Participants moved their hand in same sequence squares flashed in WM group (not in control group). (**d**) Experimental design depicting Exp-1 (NoReact-ImmediateGen, N = 32) with short training on Day-1, secondary task (WM group, N = 16; control group, N = 16) with right hand, and testing untrained left arm for immediate interlimb generalization on Day-2. Same for Exp-2 (React-ImmediateGen, N = 32) except that brief reactivation was given before secondary task (WM group, N = 16; control, N = 16), and immediate generalization was assessed immediately after secondary task. Finally, Exp-3 (React-DelayedGen, N = 31) contained short training on Day-1, brief reactivation and secondary task (WM group, N = 16; control group = 15) on Day-2 and delayed generalization was tested on Day-3 with their untrained arm.

### Task Apparatus

The experimental setup is adapted from our prior work (Yadav, Pal, et al., 2025) in which participants performed a motor skill task on a digitizing tablet (GTCO CalComp) placed beneath a horizontally mounted computer screen (1920 × 1080 pixels), with a reflective mirror positioned between the screen and the tablet. This setup prevented direct hand vision while allowing real-time visual feedback of their hand movements via a cursor on the screen. The task was programmed using Psychophysics Toolbox in MATLAB (Brainard, 1997) and hand position data were recorded at a 120 Hz sampling rate.

### Motor Skill Task

The motor skill task required participants to perform fast and accurate reaching movements between a start position and one of five target locations with the right hand holding the stylus on the digitizing tablet. The trial started with moving the hand inside the start circle. After 500 ms within the start circle (green color, 10 mm diameter), a blue target circle (10 mm diameter) appeared on the screen at 150 mm along with an audio go cue. Participants were instructed to move their hands fast and straight from the start circle to the target circle and complete the movement in 650 ms within a specified path (a parallel yellow line connecting the start circle with the target circle). Trial duration was 2 seconds, and participants received visual performance feedback and a numerical score after each trial completion. The visual feedback consisted of: (1) a red line highlighting trajectory segments outside the specified yellow path (presence of this red line indicated low spatial accuracy or high spatial error), and (2) a red square marking hand position at 650 ms, where its distance from the target circle reflected temporal error (greater distance from target position indicated lower temporal accuracy or higher temporal error). Using the spatial and temporal feedback components, we calculated a single “motor error” score (in mm), presented as additional feedback in the screen’s top corner after each trial. Spatial error was computed as the path length (mm) deviating outside the yellow path (indicated by the red line). Temporal error was the distance (mm) from the hand position at 650 ms to the target circle. Motor error represented the sum of these two values (mm). Although participants were instructed to minimize motor error through fast and accurate movements, the specific calculation method was withheld to prevent bias towards either error component. Consequently, successful trials (reaching the target within 650 ms while staying on path) yielded motor errors near zero. We used the motor error as our primary measure in this study.

### Secondary Task (Working Memory and Control Task)

Following motor skill training, participants were asked to perform one of the secondary tasks: a working memory task (WM group) or a control task (Control group). Working memory task (WM Group) is a modified version of the visuo-spatial Corsi block-tapping task used in our previous work (Yadav, Pal, et al., 2025). Here, participants are required first to move their hand-held stylus into a blue start circle, and after 750 ms, the participants hear a beep sound followed by nine white squares on the screen. A randomly ordered sequence of 3-5 squares then flashed red (750 ms per square). Participants were instructed to observe these flashing squares and remember the order and spatial location of the flashed squares. After a second beep, participants had to move their hand (shown as a yellow cursor on the screen) in the correct order within a time limit of k+1 seconds, where k was the number of flashed squares. Participants received “Trial Successful” feedback for tapping squares in the correct sequence or “Trial Unsuccessful” if the sequence was incorrect or timed out. Participants in the Control Group completed a modified WM task without the working memory requirement, and the rest of the task procedure was similar. The squares (3–5) that turned from white to red remained red (visible on screen throughout the trial), and participants were asked to tap them in any order without touching the white squares. Participants performed a total of 80 trials of this secondary task (WM or Control) with the same right arm used to train on the motor skill task, and they were required to achieve at least 70% accuracy to be included in the final analyses.

### Study Design

The study design consisted of three sets of experiments that took place over 2-3 days, involving skill training (right dominant arm), performing the secondary task (right dominant arm), no or a brief skill reactivation (right dominant arm), and finally testing for interlimb generalization (left untrained arm). Participants were randomly assigned to one of three experiments.

#### Experiment 1 (NoReact-ImmediateGen, N=32)

On Day-1, all participants trained on the motor skill task with their right hand. They performed 50 trials (consisting of five targets presented randomly). On Day-2, they performed the secondary working memory task (N = 16) or control task (N = 16) followed by immediate skill generalization testing with the left hand (50 trials).

#### Experiment 2 (React-ImmediateGen, N=32)

In Exp-2, again all participants trained on the motor skill task with the right arm (50 trials), similar to experiment 1. On Day-2, participants performed a brief reactivation session consisting of 10 trials of the motor skill task with the right arm before completing the secondary working memory (N = 16) or control task (N = 16) for 80 trials. Immediately after the secondary task, participants performed a skill generalization test with the left arm (50 trials).

#### Experiment 3 (React-DelayedGen, N=31)

On Day-1, participants trained on the motor skill task with the right arm (50 trials), similar to Exp-1 and Exp-2. On Day-2, participants performed 10 reactivation trials of the motor skill task with the right arm before performing the secondary working memory (N = 15) or control task (N = 16). Participants were tested for delayed skill generalization with the left arm (50 trials) 24-hour after the reactivation on Day-3.

### Data analysis and statistical tests

A total of ninety-five (out of 96 tested) participants were included in the analyses (one subject excluded due to a technical error during the task execution resulting in missing movement related data points). In these participants, we tested whether a secondary WM task interferes with reactivation-induced skill memory reconsolidation and impacts the follow-up interlimb generalization. To do so, we first quantified skill learning in all participants by assessing motor skill error over 50 trials of the training session. The motor error was computed as the sum of spatial and temporal errors, following procedures described in our previous work (Yadav, Pal, et al., 2025; Yadav, Vassiliadis, et al., 2025; Yadav & Mutha, 2016, 2020). This composite error value (in mm) was used to assess learning and generalization. To reduce directional bias, motor error values from five consecutive trials (corresponding to the five directional targets) were averaged to form bins (10 bins in all experiments as mentioned earlier in the experimental design).

Mixed analysis of variance (ANOVA) was used to assess learning and interlimb generalization across the three experiments, with the significance level set at 0.05. Where significant main effects were observed, pairwise t-tests were conducted and corrected for multiple comparisons using the Bonferroni method. Effect sizes were reported as omega-squared (ω^2^). The Bayesian approach further confirmed each null finding by calculating the Bayes factor in R with the library of BayesFactor at BF_10_.

Reduction in skill error across bins indicated successful learning during the training session. Learning was assessed using a two-way repeated-measures ANOVA with bin (first, last training) as a within-subject factor and group (WM, Control) as a between-subject factor, consistent with our prior study. For skill generalization from the right to the left arm, the first trial of the generalization session was excluded, consistent with the work of (Wang & Sainburg, 2006) and our previous work (Yadav, Pal, et al., 2025), who reported that the nervous system uses the initial trial to evaluate the relevance of prior learning in a new context. ANOVA was then performed with bin (first training, first generalization) as a within-subject factor and group (WM & control) as a between-subject factor, separately for each experiment.

## RESULTS

### Experiment 1: WM engagement on Day-2 does not impair interlimb generalization

In Exp-1 (NoReact-ImmediateGen), participants successfully learned the motor skill, as evidenced by a progressive reduction in error across the training session. This learning is demonstrated by the hand trajectories of a representative participant (Fig. 2a). During the initial bin (first five trials), the hand position at the 650 ms mark (indicated by a square) was close to the start circle, and movement paths were frequently deviating outside the specified path. However, trajectories from the final training bin show a marked improvement: hand positions consistently reached the target circle, and movements were straighter, remaining within the designated path. Both WM and control groups reduced errors and improved their performance over the course of training (Fig. 2b). Statistical examination revealed a significant effect of bin (first, last bin training) [*F*(1, 30) = 198.80, *p* < 0.001, ω^2^ = 0.16]. We found no main significant effect of group [*F*(1, 30) = 0.0547, *p* = 0.816, ω^2^ = 0.0012], with strong support for the null hypothesis from the Bayesian analysis (BF_10_ = 0.32) and bin × group interactions effect [*F*(1, 30) = 0.4257, *p* = 0.519, ω^2^ = 0.001] was also not significant, with strong support for the null hypothesis from the Bayesian analysis (BF_10_ = 0.37), suggesting similar amount of learning in both groups.

**Figure 2:**
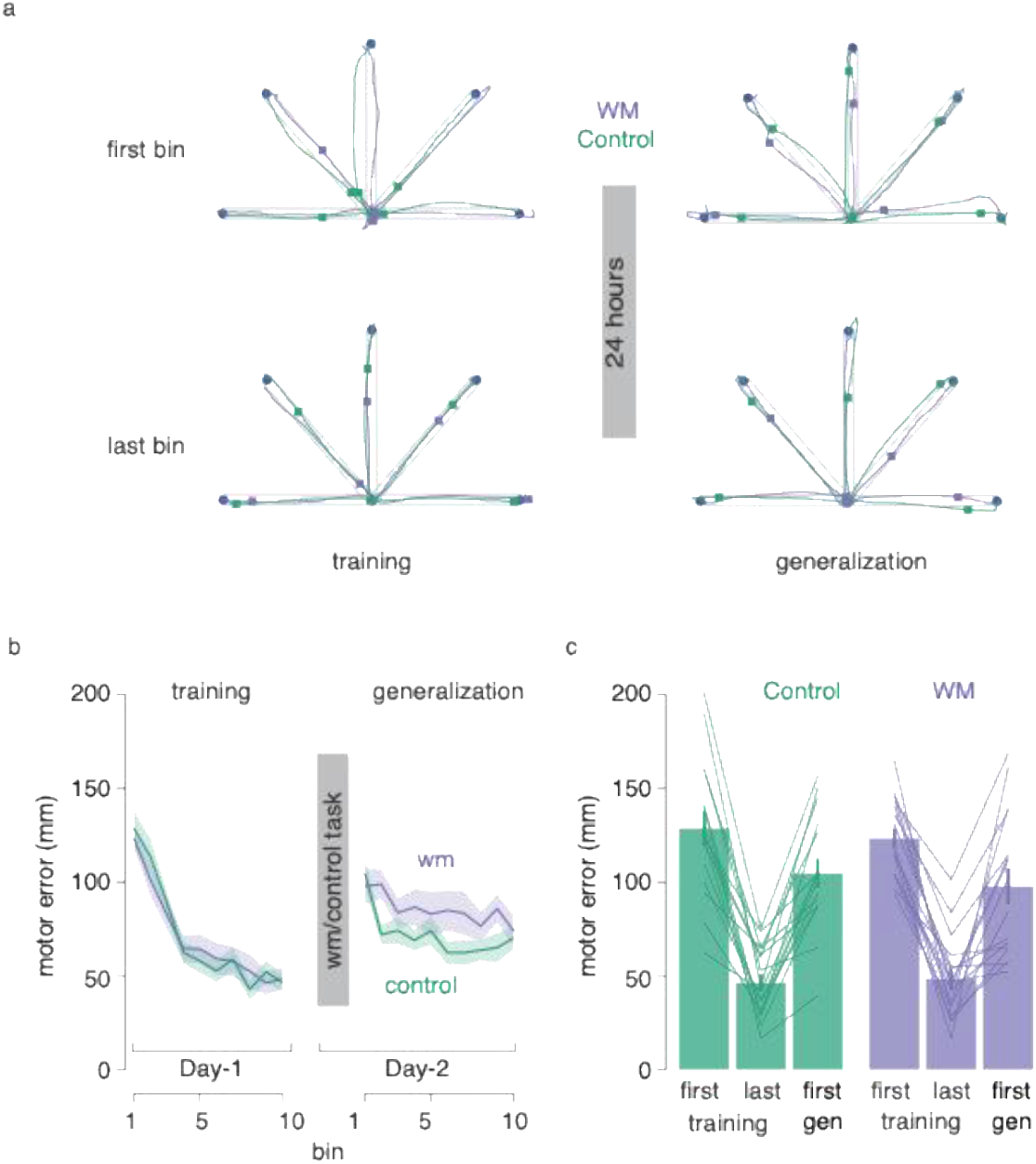
Working Memory engagement after consolidation doesn’t impair interlimb generalization. (**a**) Movement trajectories to the five targets (blue circle) of first and last bin of skill training and generalization sessions, respectively are shown for one participant from each group (WM group in purple, control in green). The position of the square (hand position at 650 ms) is closer to the target in the last bin of the training session when compared to the first bin of the training session. Similarly, it is also closer to the target in the first and last bins of the generalization session when compared to the first bin of training. This indicates improved performance by the end of training and subsequent generalization of that performance to the untrained arm in participants from both groups. (**b**) Change in mean motor error during the training and generalization sessions over the course of training for WM group (N = 16) and control group (N = 16). Shaded regions represent standard error (SE). (**c**) Bar plots comparing motor errors on first, last bins of training and first bin of generalization for the control group (green) and WM group (purple) are shown. Lines represent performance of individual participants. Error bars represent SE.

Interestingly, participants in both groups exhibited interlimb generalization of acquired motor skills to the untrained arm (lowered motor errors) when tested immediately after engaging in the secondary task. As shown in a representative participant’s trajectories (Fig. 2a), hand paths remained within the specified trajectory during the first bin of generalization, and the hand position at 650 ms consistently reached the target. We noted a significant effect of bin (first bin training, first bin generalization) [*F*(1, 30) = 12.144, *p* = 0.0015, ω^2^ = 0.009], but no group [*F*(1, 30) = 0.443, *p* = 0.510, ω^2^ = 0.0008, BF_10_ = 0.39] or bin × group interactions [*F*(1, 30) = 0.0108, *p* = 0.9178, ω^2^ = 0.001, BF_10_ = 0.33] suggesting similar levels of interlimb generalization in both WM and control groups. Critically, performance during the generalization session was not influenced by the type of secondary task performed beforehand (Fig. 2c).

Further supporting this finding, an independent samples t-test comparing the averaged motor error during the generalization session revealed no significant difference between the WM group (mean motor error = 85.51) and the control group (mean motor error = 71.84), [*t*_*[30]*_ = 1.40, *p* = 0.173, *Cohen’s d* = 0.494].

### Experiment 2: WM engagement after skill reactivation on Day-2 enhances immediate interlimb generalization

In Exp-2 (React-ImmediateGen) also participants were exposed to the similar short training on Day-1. Participants successfully reduced the error (see hand paths of a representative participant in Fig. 3a, and group data in Fig. 3b) during the course of training with a significant effect of bin (first bin training, last bin training) [*F*[1, 30] = 196.71, *p*<0.001, ω2 = 0.1524] but no group [*F*[1, 30] = 1.36, *p* = 0.25, ω2 = 0.00066, BF_10_ = 0.57] or bin X group interaction [*F*[1, 30] = 0.671, *p* = 0.419, ω2 = 0.0012, BF_10_ = 0.44]. This suggests that participants from both groups acquired similar skill performance.

**Figure 3:**
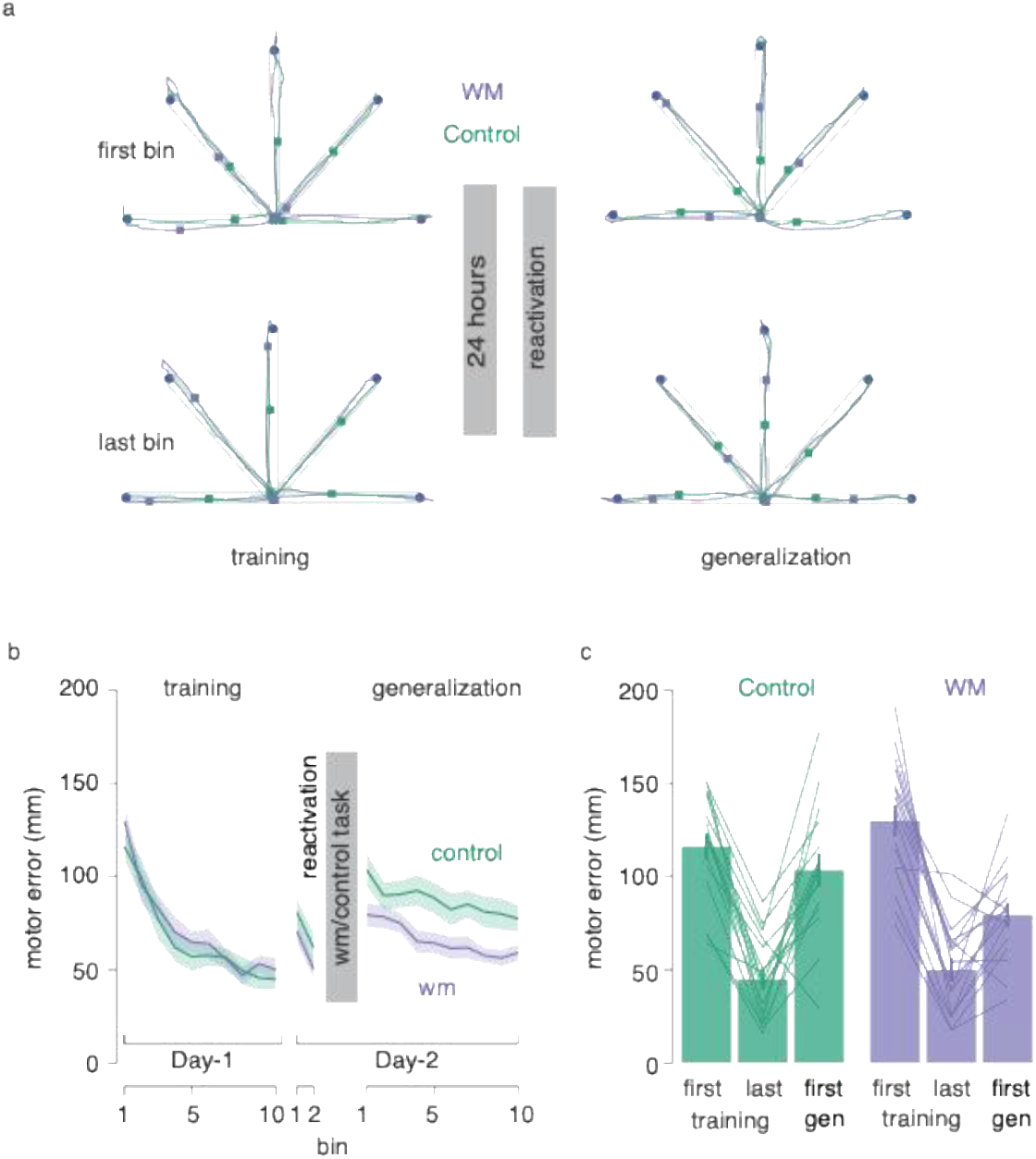
Working memory engagement after memory reactivation enhances immediate interlimb generalization. (**a**) Movement trajectories to the five targets (blue circle) of first and last bin of skill training and generalization sessions, respectively are shown for one participant from each group (working memory in purple, control in green). Here, as compared to the first bin of training, position of square (hand position at 650 ms) is close to the target in last bin of training and in the first and last bins of generalization; indicating improved learning over the course of training and subsequent generalization of that performance to the untrained arm in participants of both groups. (**b**) Change in mean motor error during training and generalization sessions over bins for WM (N = 16) and control group (N = 16). Shaded regions represent standard error (SE). (**c**) Bar plots comparing motor errors in the first and last bins of training, as well as the first bin of generalization, are shown for the control group (green) and the WM group (purple). Lines represent the performance of individual participants. The error bars represent SE.

Participants were exposed to the reactivation session on Day-2 after consolidation of initial training. Participants successfully reduced their error (Fig. 3b), as indicated by a significant main effect of bin (first bin reactivation, last bin reactivation) [*F*(1, 30) = 35.59, *p* < 0.001, ω^2^ = 0.015], and both groups performed similar [*F*(1, 30) = 1.33, *p* = 0.257, ω^2^ = 0.0014, BF_10_ = 0.63]. Further, by the end of reactivation, both groups have reached the same level of performance as the end of Day-1 training [*F*(1, 30) = 2.40, *p* = 0.132, ω^2^ = 0.0012].

Next and central to our study, interlimb generalization was evident when participants were tested for generalization with their left arm after 24-hour of training. Hand paths were within the specified trajectory during the first bin of generalization, and the hand position at 650 ms consistently reached the target (Fig. 3a). We found a significant interaction between bin and group [*F*(1, 30) = 7.690, *p* = 0.00945, ω^2^ = 0.0058]. Posthoc analysis revealed that WM group showed enhanced performance in interlimb generalization (*p* = 0.034) than control group (Fig. 3c). Further, there was a significant difference in motor error (averaged bin) between WM (mean motor error = 65.87 SE) and control group (mean motor error = 87.06 SE), [*t*_*[30]*_ = -2.69, *p* = 0.01155, *Cohen’s d* = -0.951] which suggest that the enhanced interlimb generalization was maintained throughout the generalization session

### Experiment 3: Skill reactivation followed by WM engagement and 24-hour reconsolidation window leads to impaired interlimb generalization on Day-3

In Exp-3 (React-DelayedGen), participants successfully learned the skill over the course of the training session (see Fig. 4a for hand trajectories of a representative participant, and Fig. 4b for the group data) with a significant effect of bin (first, last bin training) [*F*(1, 29) = 202.199, *p*<0.001, ω^2^ = 0.164]. There was no significant main effect of group [*F*(1, 29) = 0.0172, *p* = 0.8963, ω^2^ = 0.0014, BF_10_ = 0.34] or interaction between bin × group [*F*(1, 29) = 0.2298, *p* = 0.6352, ω^2^ = 0.0013, BF_10_ = 0.38].

**Figure 4:**
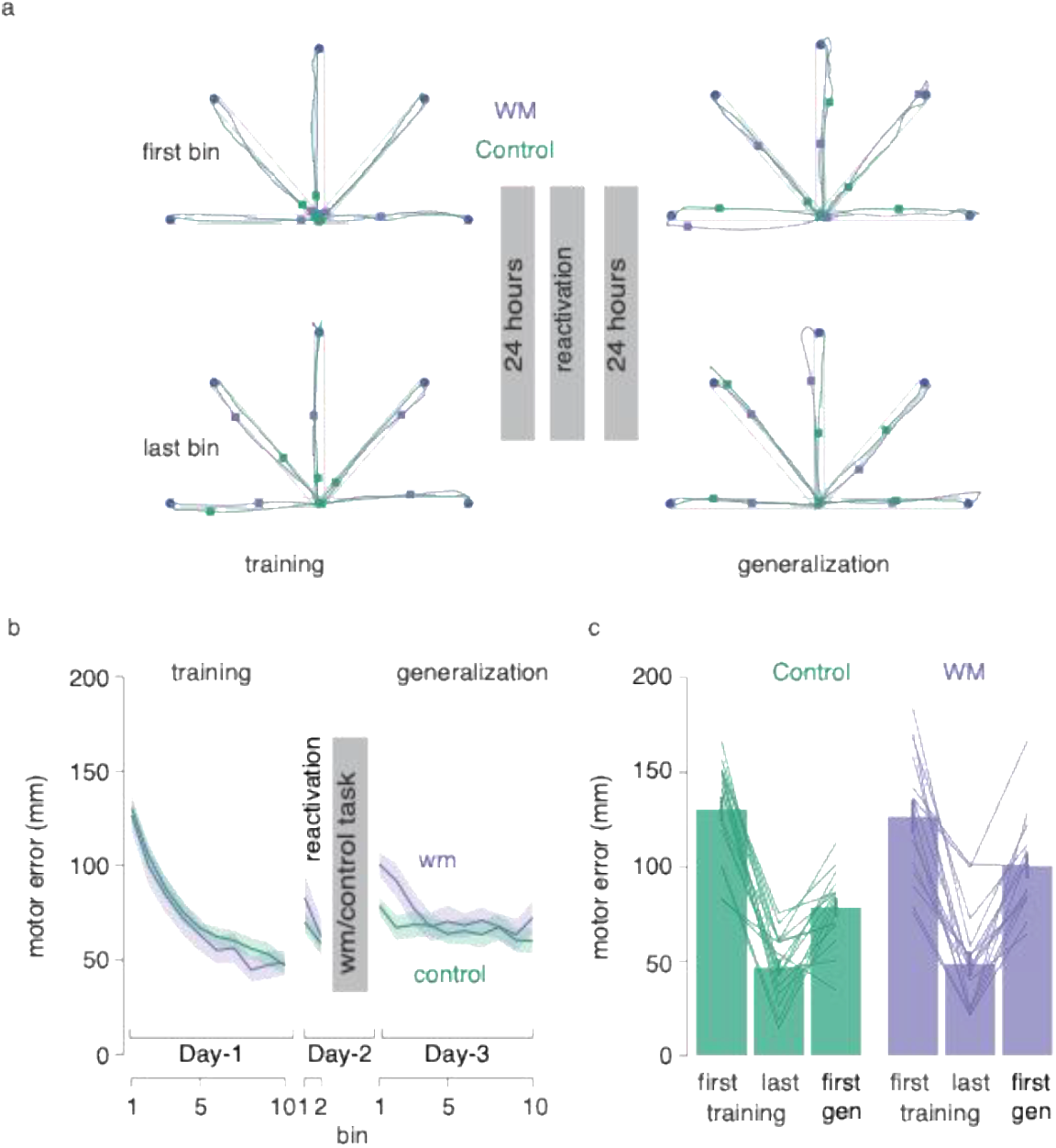
Working memory engagement after memory reactivation enhances delayed interlimb generalization. (**a**) Movement trajectories to the five targets (blue circle) of first and last bin of skill training and generalization sessions, respectively are shown for one participant from each group (working memory in purple, control in green). Similar to Exp-1 and Exp-2, as compared to the first bin of training, position of square (hand position at 650 ms) is close to the target in last bin of training and the first and last bins of generalization, indicating improved learning over the course of training and subsequent generalization of that performance to the untrained arm in the participants of both groups. (**b**) Change in mean motor error during training and generalization sessions for WM group (N = 15) and control group (N = 16). Shaded regions represent standard error (SE). (**c**) Bar plots comparing motor errors in the first and last bins of training, as well as the first bin of generalization, are shown for the control group (green) and the WM group (purple). Lines represent the performance of individual participants, and error bars indicate standard error (SE).

Similar to Exp-2, during the reactivation session, participants showed a significant reduction in error [*F*(1, 29) = 16.93, *p* = 0.0003, ω^2^ = 0.0051] over the two bins, and there was no significant effect of group [*F*(1, 29) = 0.50, *p* = 0.485, ω^2^ = –0.003, BF_10_ = 0.62]. Further, by the end of reactivation, both groups have reached the same level of performance as the end of day-1 training [*F*(1, 29) = 0.02, *p* = 0.887, ω^2^ = –0.0040].

Finally, participants showed interlimb generalization when tested on Day-3. Fig. 4a shows the hand movement within the specified path and the hand position at 650 ms consistently reached the target (Fig. 4a). We noted a significant bin × group interaction effect [*F*(1, 29) = 4.766, *p* = 0.0372, ω^2^ = 0.02], suggesting that the degree of interlimb generalization differed between both groups. Post-hoc test revealed that the interlimb generalization of participants in WM group (mean motor error = 100.58) was impaired than those in the control group (mean motor error = 78.68) (*p* = 0.011). However, this impairment was not long-lasting and there was no significant difference in mean motor error between WM (mean motor error = 75.09) and control group (mean motor error = 66.33), [*t*_*[29]*_ = 1.00, *p* = 0.3259, *Cohen’s d* = 0.3593] across the generalization session.

Furthermore, to investigate the influence of reactivation on interaction between WM and interlimb generalization, we pooled and directly compared the performance of WM groups during the generalization session across Exp-2 & Exp-3. In Exp-3, participants underwent a reconsolidation (restabilization) process characterized by a 24-hour gap between the reactivation session (followed by a secondary task) and the subsequent generalization session. In contrast, participants in Exp-2 completed the generalization session immediately following the reactivation and secondary task intervention on the same day. We combined WM participants from both Exp-2 & Exp-3 and conducted two-sample independent t-test between first bin of generalization. The results revealed a statistically significant difference between the groups, [*t*_*[29]*_ = -2.31, *p* = 0.028, *Cohen’s d* = 0.83]. Specifically, participants in the WM group from Exp-3 (mean motor error = 100.58), who underwent a 24-hour reconsolidation period, showed impaired generalization compared to those in the WM group from Exp-2 (mean motor error = 79.42), who were tested while the memory was still in a labile state. Similarly, we combined control group participants from both Exp-2 and Exp-3 and compared the first bin of generalization. The results revealed a significant difference between the experiments, [*t*_*[30]*_ = 2.46, *p* = 0.020, *Cohen’s d* = 0.87]. Participants in control group from Exp-2 (mean motor error = 103.62) exhibited significantly higher error scores than those in control group from Exp-3 (mean motor error = 78.68).

## DISCUSSION

In the present study, we investigated how engagement of working memory following skill memory reactivation influences interlimb generalization. Building on our previous finding that WM engagement following a short skill training impairs delayed generalization (Yadav, Pal, et al., 2025), we hypothesized that WM engagement after a brief skill reactivation would similarly disrupt generalization to the untrained arm. However, our results revealed a more nuanced pattern-WM engagement following reactivation at 24-hour post-training enhanced generalization when tested immediately (Exp-2) but impaired generalization when tested after an additional 24-hour delay (Exp-3). These seemingly contradictory findings can be reconciled by a unified framework in which WM engagement prolongs the labile state of the reactivated skill memory which is beneficial for immediate transfer but detrimental with a time gap.

### The necessity of skill reactivation skill memory-WM interaction following skill stabilization/consolidation

In Exp-1, with no skill reactivation, WM engagement on Day-2 did not affect interlimb generalization compared to the control group. This finding establishes a critical boundary condition: a consolidated motor memory, once stabilized, is resistant to interference from a secondary WM task (Brashers-Krug et al., 1996; Robertson, 2018). Without reactivation, the skill memory trace remains in its stable state from Day-1 training, therefore unperturbed by WM engagement at 24-hour. This result aligns with the reconsolidation literature, which posits that memory lability must be reinstated through retrieval or brief re-exposure to allow modification (Alberini & LeDoux, 2013; Nader et al., 2000; Nader & Hardt, 2009). Thus, skill memory reactivation interestingly serves as the gateway for subsequent WM engagement to influence the skill memory trace. We demonstrate this with Exp-2, wherein participants received a brief reactivation prior to performing the WM/control task on Day-2. Here, contrary to our initial hypothesis, we observed greater interlimb generalization for WM group as compared to the control group suggesting that WM engagement does not simply disrupt the reactivated memory but instead actively shapes it in a manner that facilitates transfer. We propose that the WM task prolongs the labile state of the reactivated skill memory. The cognitively demanding WM task, which engages dorsolateral prefrontal cortex (DLPFC) and other executive networks common to skill learning (Curtis & D’Esposito, 2003; Miller & Cohen, 2001), may sustain the activation of these neural circuits, effectively extending the duration of this labile window for the skill memory. Every memory representation has both general/abstract characteristics that apply to all contexts, and specific characteristics that apply only to one context. For successful generalization, the abstract characteristics are required in new contexts. Critically, a memory in an unstable/labile state is characterized by greater flexibility and accessibility of its abstract, effector-independent components (Mosha & Robertson, 2016). When generalization is tested immediately, the skill memory which is in the extended unstable phase is readily transferable to the untrained left arm, resulting in enhanced generalization performance.

The control group’s performance in Exp-2 provides convergent evidence for this interpretation. Without the cognitive demands of the WM task, the reactivated memory likely began its natural re-stabilization process rapidly. This rapid re-stabilization may favor the formation of effector-specific representations optimized for the trained right arm, leaving the memory less accessible for transfer to the untrained limb when tested immediately after brief reactivation and control task sessions.

### Time window following reactivation critical for skill memory-WM interaction

In Exp-3, the identical intervention (reactivation followed by WM task on Day-2) led to impaired generalization when tested following an additional gap of 24-hour. This delayed impairment effect mirrors our previous finding where WM engagement immediately after training disrupted transfer (Yadav, Pal, et al., 2025), suggesting that consolidation of post-training memory and reconsolidation of a future occurrence of the original memory share common underlying mechanisms.

During the 24-hour interval between the reactivation and the generalization test, the reactivated memory must undergo active re-stabilization - a process of reconsolidation that requires protein synthesis, synaptic reorganization, and systems-level consolidation (De Beukelaar et al., 2016; Dudai, 2012; Pedreira et al., 2002). Prior engagement with the WM task may create a resource competition that impairs such reconsolidation process. Consequently, memory is restabilized inefficiently, leading to a corrupted or incomplete memory trace that fails to support robust generalization to the untrained arm. The control group in Exp-3 demonstrates what successful reconsolidation looks like. Without resource competition from the WM task, the reactivated memory undergoes complete, undisturbed re-stabilization. Notably, this 24-hour re-stabilization period, potentially involving sleep-dependent processes (Cohen et al., 2005; Walker et al., 2003), may strengthen the memory’s abstract, generalizable components, leading to robust delayed generalization superior to the immediate generalization observed in the Exp-2 control group.

The apparent paradox of WM enhancing immediate generalization but impairing delayed generalization can be resolved by considering the temporal dynamics of the reconsolidation window. We propose that reconsolidation comprises at least two distinct phases: an initial period of lability and malleability, followed by a subsequent period of active re-stabilization that requires dedicated neural and cognitive resources. WM engagement following reactivation prolongs the initial labile phase, which is beneficial for immediate transfer but comes at the cost of disrupting the subsequent re-stabilization phase through resource competition. This framework parsimoniously explains why the same intervention produces opposite outcomes depending on when generalization is tested.

This dual effect aligns with emerging views that memory reconsolidation is not a unitary process but a dynamic sequence of events that can be modulated at different stages (Alberini & LeDoux, 2013; Dudai, 2012). Our findings also resonate with studies showing that post-reactivation manipulations can either strengthen or weaken memories depending on the nature and timing of the intervention (Amar-Halpert et al., 2017; Herszage & Censor, 2017; Wymbs et al., 2016). Critically, we extend this literature by demonstrating that a single cognitive intervention can produce bidirectional effects on a motor memory based solely on when the memory is probed.

### Implications and Future Directions

Our findings have important implications for understanding the functional architecture of memory systems. First, we provided compelling evidence for cross-system interactions between declarative (WM) and procedural (motor skill) memory systems during reconsolidation. These interactions are not simply competitive but can be facilitatory depending on timing (Brown & Robertson, 2007; Robertson, 2018). Second, our results suggest that the neural substrates of reconsolidation, including DLPFC and related circuits, may be shared resources that can be co-opted by concurrent cognitive demands, with consequences for memory integrity. From a translational perspective, these findings may inform neurorehabilitation strategies. Interventions that combine skill reactivation with cognitive engagement could be timed to either promote immediate flexibility (e.g., for adapting to novel contexts) or ensure long-term retention (e.g., for permanent skill acquisition). Understanding how to optimize this temporal window could enhance the efficacy of motor rehabilitation protocols. Future research should directly investigate the neural mechanisms underlying these effects. Neuroimaging or non-invasive brain stimulation studies could test whether DLPFC engagement during the reactivation-WM intervention predicts both immediate facilitation and delayed impairment. Additionally, manipulating the duration and timing of reactivation as well as the re-stabilization window could further elucidate the critical time period during which resource competition exerts its disruptive effects.

## Acknowledgments

We would like to thank Dr. Leonardo G. Cohen for his valuable feedback on the manuscript.

